# Sampling bias in large healthcare claims databases

**DOI:** 10.1101/2022.10.03.510721

**Authors:** Alex Dahlen, Vivek Charu

## Abstract

Healthcare claims databases that aggregate claims from multiple commercial insurers are increasingly being used to generate real-world evidence. These databases represent a non-random sample of the underlying population, but often little attention is paid to the inherent sampling bias within the data, and how it might affect results. As an illustrative example, we characterize variation in sampling in Optum's de-identified Clinformatics Data Mart Database (CDM) at the zip-code level in 2018, and identify socioeconomic and demographic factors associated with inclusion.

## Introduction

Healthcare claims databases that aggregate claims from multiple commercial insurers are increasingly being used to generate real-world evidence^1–4^. These databases represent a non-random sample of the underlying population, but often little attention is paid to the inherent sampling bias within the data, and how it might affect results. As an illustrative example, we characterize variation in sampling in Optum’s de-identified Clinformatics Data Mart Database (CDM) at the zip-code level in 2018, and identify socioeconomic and demographic factors associated with inclusion.

## Methods

Optum’s CDM consists of administrative claims derived from several large commercial and Medicare Advantage health plans. For our primary analysis, we count the number of individuals with CDM coverage on a single day (June 1, 2018) in each zip code, and compare it to the 2018 census estimates of the total population in that zip code. In sensitivity analyses, we also consider one strictly larger cohort (individuals with at least one day of coverage at any point during 2018) and one strictly smaller cohort (individuals with continuous coverage during the entire year of 2018).

To explain the variation in zip-code level CDM sampling, we fit an inverse-variance weighted multivariable linear regression model with 29 socioeconomic and demographic features extracted from the 2018 census, and state-level fixed effects. See **Table 1** for the list of features, and the **Appendix 1** for more details about the model and methods.

**Table 1.** Associations between 29 socioeconomic and demographic features and CDM sampling fraction at the zip-code level, accounting for state-level variation in sampling fraction in two models. The first set of models considers each covariate of interest separately, along with state-level fixed effects. Partial correlation coefficients derived from this model are presented; positive correlations are colored in red, and indicate that zip codes that with higher values of the covariate of interest are associated with higher zip-code level sampling in CDM, even after adjusting for state-level clustering in sampling. The second model is a full multivariable model that includes all 29 covariates of interest in addition to state-level fixed effects. For example, for a 10 percentage increase in a zip code’s fraction of households earning greater than $200,000, the model suggests the CDM sampling fraction will increase by 0.6 percentage points, on average. Full model results (including state-level fixed effects) are provided in **Table 3** below.

|  |  | Partial Corr.<br>Coeff. | Multivariable Regression<br>Coefficient | p |
| --- | --- | --- | --- | --- |
| Population | Total Population (millions) | 0.01 | -0.0005 ± 0.00018 | < 0.001 |
|  | Log<sub>10</sub> Pop. Density (1/sq. mile) | 0.14 | 0.67 ± 0.06 | < 0.001 |
| Gender | % Female | 0.12 | 0.050 ± 0.010 | < 0.001 |
| Race/Ethnicity | % White | 0.20 | Comparison |  |
|  | % Black | -0.15 | -0.008 ± 0.002 | < 0.001 |
|  | % Hispanic | -0.19 | 0.001 ± 0.004 | 0.64 |
|  | % Asian | 0.16 | -0.010 ± 0.004 | < 0.001 |
|  | % Other | -0.08 | -0.026 ± 0.008 | < 0.001 |
| Age | % Under 18 | -0.09 | Comparison |  |
|  | % 18 - 40 | -0.21 | -0.019 ± 0.010 | < 0.001 |
|  | % 40 - 60 | 0.32 | 0.084 ± 0.014 | < 0.001 |
|  | % 60 - 80 | 0.14 | 0.002 ± 0.012 | 0.71 |
|  | % Over 80 | 0.12 | 0.055 ± 0.024 | < 0.001 |
| Household<br>income | % < \$15k | -0.34 | Comparison | |
| | % \$15k – \$30k | -0.40 | -0.016 ± 0.014 | 0.014 |
| | % \$30k – \$45k | -0.37 | -0.009 ± 0.012 | 0.16 |
| | % \$45k – \$60k | -0.23 | 0.002 ± 0.014 | 0.78 |
| | % \$60k – \$100k | 0.10 | 0.007 ± 0.010 | 0.15 |
| | % \$100k – \$125k | 0.33 | 0.033 ± 0.018 | < 0.001 |
| | % \$125k – \$200k | 0.43 | 0.016 ± 0.012 | 0.011 |
| | % > \$200k | 0.42 | 0.071 ± 0.014 | < 0.001 |
| Work &<br>Insurance | % Unemployed | -0.29 | -0.001 ± 0.014 | 0.89 |
|  | % No health insurance | -0.31 | -0.030 ± 0.008 | < 0.001 |
| Education | % Less than High School | -0.33 | Comparison |  |
|  | % High School | -0.27 | 0.038 ± 0.01 | < 0.001 |
|  | % Some College | -0.09 | -0.010 ± 0.008 | 0.014 |
|  | % College | 0.41 | 0.071 ± 0.010 | < 0.001 |
|  | % Graduate | 0.30 | -0.021 ± 0.010 | < 0.001 |
| Housing | % houses that are owner occupied | 0.25 | -0.003 ± 0.004 | 0.14 |
| | Median house price (millions of \$) | 0.36 | -0.00001 ± 0.00003 | 0.43 |

## Results

16.4 million distinct individuals were captured in CDM on June 1, 2018, representing 5.4% of the US population. The median zip code sampling fraction was 4.4% (interquartile range: 2.5-7.1%), with clear geographic variation in sampling (**Figure 1**). At the state level, Alaska had the lowest sampling rate (0.9%) and Colorado had the highest sampling rate (10.9%); in multivariable regression models, state-level fixed effects explained 34.6% of the zip-code level variation in sampling.

**Figure 1.**
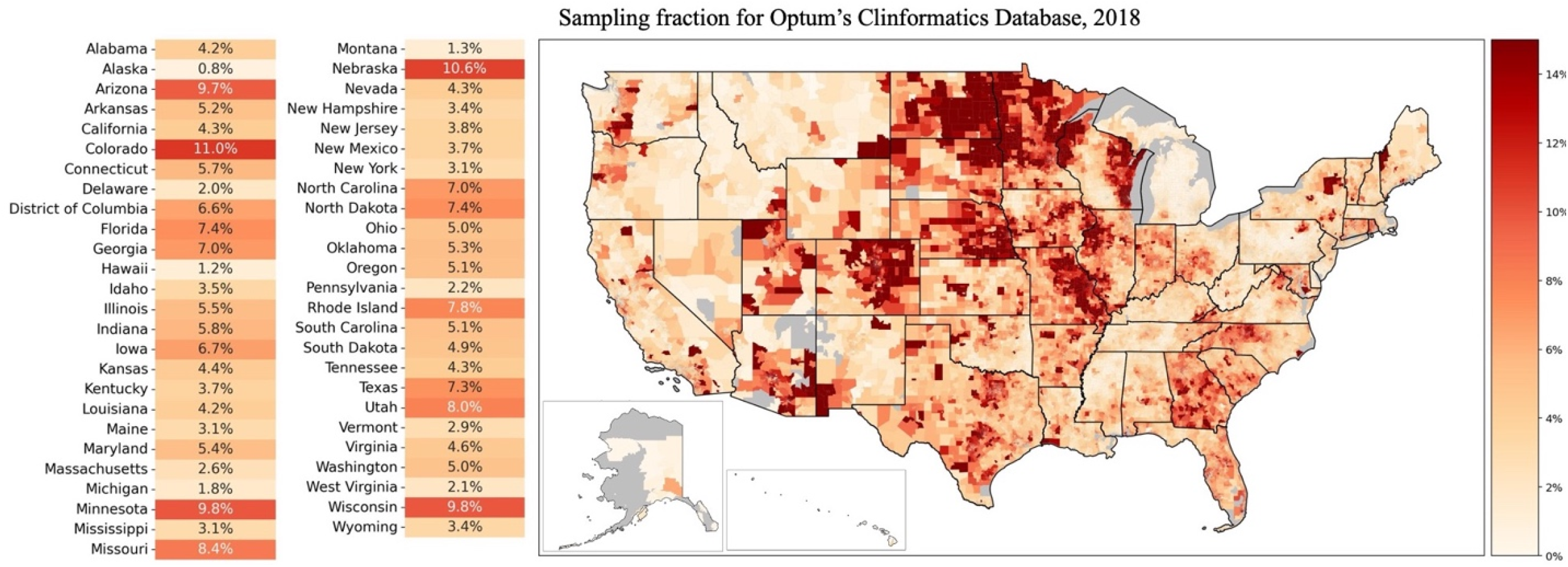
Zip-code and state-level variation in sampling in Optum’s Clinformatics® Data Mart Database (CDM) in 2018. The right panel displays CDM sampling in each zip-code, estimated from the number of patients with coverage in CDM on June 1, 2018. The left panel displays the estimated state-level sampling using the same criteria.

Associations between socioeconomic/demographic features and CDM sampling fraction, after adjusting for state-level variation, are provided in **Table 1**. Estimated partial correlations and regression model coefficients demonstrate that inclusion in CDM is associated with zip codes that are wealthier, older, more educated, and disproportionately White. These patterns are robust to the choice of cohort definition, and across the 10 most populous states individually (**Appendix 2**) Socioeconomic and demographic features explain an additional 19.4% of the zip code level-variation in sampling on top of state-level variation, for a total adjusted R^2^ of 54.0%.

## Discussion

To interpret results generated from healthcare claims databases, it is essential understand which patients are represented in them. We demonstrate that inclusion in Optum’s CDM at the zip-code level in 2018 varies spatially and along socioeconomic and demographic lines. We analyzed data at the smallest geographic scale available in CDM, the zip code; given our findings, there is likely to be additional bias within zip codes as well.

The same socioeconomic and demographic features that correlate with over-representation in claims data have also been shown to be effect modifiers across a diverse spectrum of health outcomes, including the burden of disease and access to healthcare^5-6^. This combination of heterogenous sampling and effect modification—both driven, in this case, by social determinants of health—gives rise to external validity bias, where results generated from the claims data will fail to generalize to the underlying population^7-8^. This external validity bias can affect both studies that estimate disease incidence or prevalence and comparative effectiveness studies that use contemporary causal inference methods: when there is meaningful heterogeneity in treatment/policy effects along socioeconomic or demographic lines, then claims-derived estimates will be biased.

Our study highlights the importance of investigating sampling heterogeneity in analyses of large healthcare claims data to evaluate how sampling bias might compromise the accuracy and generalizability of results^9^. Importantly, investigating these biases or accurately correcting for such bias will require external data sources outside of the claims database itself. Healthcare claims databases offer enormous promise for medical research; characterizing and overcoming sampling bias in these datasets is essential.

## Acknowledgement

We thank Kate Miller and Steve Goodman for conversations about the manuscript. Data for this project were accessed using Stanford Center for Population Health Sciences (PHS) Data Core. The PHS Data Core is supported by a National Institutes of Health National Center for Advancing Translational Science Clinical and Translational Science Award (UL1TR003142) and from Internal Stanford funding. The content is solely the responsibility of the authors and does not necessarily represent the official views of the NIH. Dataset DOI: 10.57761/y4af-nd44.

## Appendix 1: Detailed Methods

Here we describe additional details about the methodology used.

Optum’s de-identified Clinformatics® Data Mart Database (CDM) is a de-identified database derived from a large adjudicated claims data warehouse. We considered three cohort definitions: (1) all unique patients in CDM on June 1, 2018; (2) all unique patients with at least a single day of coverage from January 1, 2018 to December 31, 2018 and (3) all unique patients with coverage during the entire year, January 1, 2018 to December 31, 2018. Cohort 1 parallels a common use of the that computes incidence or prevalence rates by normalizing counts by patient-days of coverage; cohort 2 is the maximal count of covered patients in 2018; and cohort 3 parallels a different common use of the data where cohorts with longer periods of continuous coverage are compared.

For each cohort definition, we estimated the zip-code level CDM sampling fraction as the number of unique patients in each zip code in CDM divided by the number of unique persons in the American Community Survey (ACS) 2018 5-year census. Of note, CDM aggregates some smaller zip-codes into groups; all analyses are performed at the level of CDM’s zip-code clusters definition.

Socioeconomic/demographic information was derived from the ACS 2018 5-year census, and included population density; the fraction of persons identifying as Black, Asian, White, Hispanic, or other races; the fraction of persons aged <18, 18-39, 40-59, 60-19, and >80; the fraction of households earning (in US dollars) less than $15,000, $15,000-$30,000, $30,000-$45,000, $45,000-$60,000, $60,000-$100,000, $100,000-$125,000, $125,000-$200,000, and over $200,000; the fraction of individuals unemployed and the fraction of individuals without health insurance; the fraction of individuals with who completed less than high school, high school, some college, college, and graduate school; the fraction of houses owner-occupied; and the median house price. This data was read in at the census track level and then to get the data on equal footing, we rolled the census data up using a population-weighting or square mileage-weighting, as appropriate.

The multivariable model is a linear regression of the form:

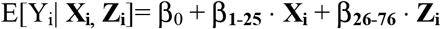

where *i* indexes each zip-code, Y_i_ is the CDM sampling fraction in each zip code, **X**_**i**_ represents a vector of 25 census features for each zip code (29 total socioeconomic and demographic features with 4 comparisons groups) and **Z**_**i**_ represents a vector of 50 state-level fixed effects (51 total states including the District of Columbia with one comparison group). Each zip code is weighted by the inverse of the variance associated with the zip code level estimate of CDM sampling. We used the standard binomial estimate of variance, modified by Winsorizing the estimate of the sampling fraction at the 5^th^ percentile, to prevent unstable weighting. Thus, the weights used scale with the total census population in each zip-code, down-weighting the contribution of very small and high variance zip codes. We produced standard diagnostic plots for this model, which demonstrated a reasonable fit.

Partial correlation coefficients were computed using the analogous short model with only a single census variable, in addition to the 50 fixed effects. The analysis was carried out in Python version 3.8.5, and the code to reproduce this analysis is publicly available at: https://github.com/alex-dahlen/who_is_in_optum.

## Appendix 2: Results of Sensitivity Analyses

Figure 3. provides unadjusted zip-code level correlation coefficients between socioeconomic and demographic features and CDM sampling for the 10 most populous states. **Table 2** provides results of sensitivity analyses in which we vary the cohort definition, as described above.

**Figure 3.**
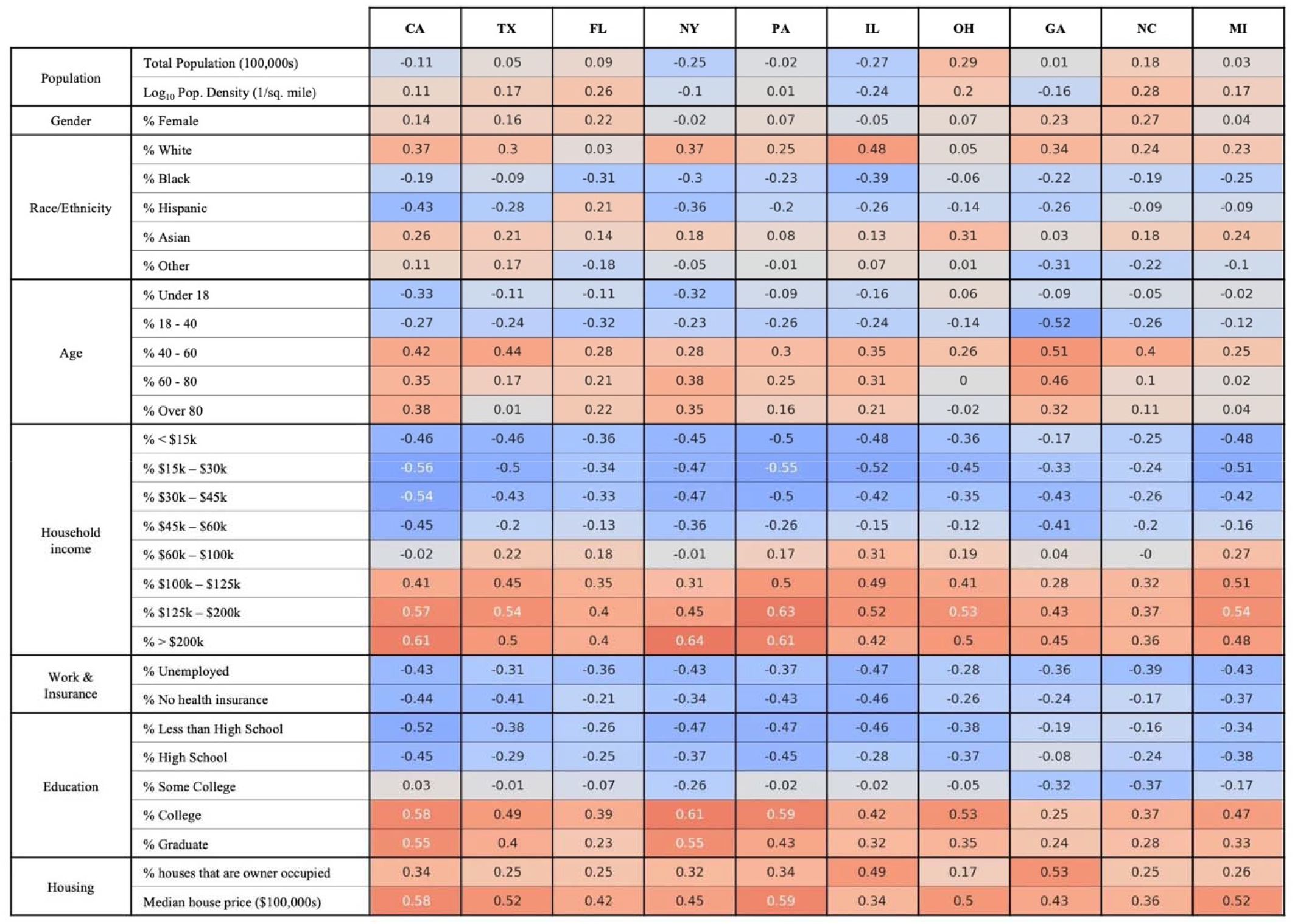
Zip code-level correlation coefficients (unadjusted and inverse variance-weighted) between the 29 sociodemographic features and CDM sampling for the 10 most populous states. The partial correlation coefficients considered in **Table 1** are derived from a model with state-level fixed effects, and thus represent an average over each state; here we consider the 10 most populous states individually. While there is some state-level variation, patterns of associations identified are consistent with the results presented in the main text.

**Table 2.** The results of two sensitivity analyses varying the cohort definition. Our primary analysis considered all patients with CDM coverage on June 1, 2018. Here we consider: all patients with at least one day of coverage at any point during the entirety of 2018; and all patients who are covered for the entire year of 2018. Results are robust to these changes in cohort definition.

|  |  | Covered for at least 1 day in 2018 |  |  | Covered for the entirety of 2018 |  |  |
| --- | --- | --- | --- | --- | --- | --- | --- |
|  |  | Partial Corr. Coeff. | Multivariable Regression Coefficient | p | Partial Corr. Coeff. | Multivariable Regression Coefficient | p |
| Population | Total Population (millions) | 0.02 | -0.0007 ± 0.0001 | < 0.001 | 0.00 | -0.0003 ± 0.00014 | < 0.001 |
|  | Log<sub>10</sub> Pop. Density (1/sq. mile) | 0.16 | 0.83 ± 0.07 | < 0.001 | 0.11 | 0.51 ± 0.04 | < 0.001 |
| Gender | % Female | 0.11 | 0.07 ± 0.02 | < 0.001 | 0.12 | 0.03 ± 0.01 | < 0.001 |
| Race/Ethnicity | % White | 0.19 | Comparison |  | 0.21 | Comparison |  |
|  | % Black | -0.14 | -0.010 ± 0.002 | < 0.001 | -0.15 | -0.006 ± 0.002 | < 0.001 |
|  | % Hispanic | -0.19 | 0.000 ± 0.004 | 0.88 | -0.18 | 0.002 ± 0.002 | 0.087 |
|  | % Asian | 0.17 | -0.012 ± 0.006 | < 0.001 | 0.14 | -0.005 ± 0.004 | 0.004 |
|  | % Other | -0.07 | -0.032 ± 0.010 | < 0.001 | -0.08 | -0.020 ± 0.006 | < 0.001 |
| Age | % Under 18 | -0.09 | Comparison |  | -0.08 | Comparison |  |
|  | % 18 - 40 | -0.18 | -0.019 ± 0.012 | 0.001 | -0.24 | -0.017 ± 0.008 | < 0.001 |
|  | % 40 - 60 | 0.31 | 0.116 ± 0.018 | < 0.001 | 0.32 | 0.054 ± 0.010 | < 0.001 |
|  | % 60 - 80 | 0.12 | 0.000 ± 0.014 | 0.97 | 0.18 | 0.006 ± 0.008 | 0.19 |
|  | % Over 80 | 0.10 | 0.053 ± 0.030 | < 0.001 | 0.15 | 0.055 ± 0.018 | < 0.001 |
| Household income | % < \$15k | -0.34 | Comparison | | -0.33 | Comparison | |
| | % \$15k – \$30k | -0.41 | -0.018 ± 0.016 | 0.031 | -0.38 | -0.014 ± 0.010 | 0.005 |
| | % \$30k – \$45k | -0.38 | -0.011 ± 0.016 | 0.19 | -0.35 | -0.007 ± 0.010 | 0.14 |
| | % \$45k – \$60k | -0.24 | 0.010 ± 0.018 | 0.26 | -0.21 | -0.004 ± 0.010 | 0.52 |
| | % \$60k – \$100k | 0.09 | 0.018 ± 0.012 | 0.003 | 0.10 | -0.001 ± 0.008 | 0.72 |
| | % \$100k – \$125k | 0.34 | 0.051 ± 0.022 | < 0.001 | 0.32 | 0.014 ± 0.014 | 0.032 |
| | % \$125k – \$200k | 0.44 | 0.024 ± 0.016 | 0.002 | 0.42 | 0.011 ± 0.010 | 0.029 |
| | % > \$200k | 0.43 | 0.098 ± 0.018 | < 0.001 | 0.40 | 0.046 ± 0.010 | < 0.001 |
| Work & Insurance | % Unemployed | -0.30 | 0.002 ± 0.016 | 0.83 | -0.28 | -0.002 ± 0.010 | 0.72 |
|  | % No health insurance | -0.31 | -0.037 ± 0.01 | < 0.001 | -0.29 | -0.021 ± 0.006 | < 0.001 |
| Education | % Less than High School | -0.33 | Comparison |  | -0.31 | Comparison |  |
|  | % High School | -0.29 | 0.050 ± 0.012 | < 0.001 | -0.23 | 0.028 ± 0.008 | < 0.001 |
|  | % Some College | -0.10 | -0.015 ± 0.010 | 0.004 | -0.07 | -0.003 ± 0.006 | 0.40 |
|  | % College | 0.43 | 0.095 ± 0.014 | < 0.001 | 0.37 | 0.05 ± 0.008 | < 0.001 |
|  | % Graduate | 0.32 | -0.026 ± 0.014 | < 0.001 | 0.27 | -0.012 ± 0.008 | 0.004 |
| Housing | % houses that are owner occupied | 0.22 | -0.012 ± 0.004 | < 0.001 | 0.27 | 0.004 ± 0.002 | 0.005 |
| | Median house price (millions of \$) | 0.37 | -0.00002 ± 0.00004 | 0.24 | 0.34 | 0.00001 ± 0.00003 | 0.94 |

## Appendix 3: Full Model Parameters

**Table 3** provides estimated parameters for the full model in the primary analysis, including the state-level fixed effects.

**Table 3.** Full model results for the multivariable regression presented in main text **Table 1**.

|  | Coef. | Std.Err. | P>\|t\| |
| --- | --- | --- | --- |
| **Intercept** | -0.0354 | 0.007 | <0.001 |
| **Total Pop (millions)** | -0.0005 | 0.0001 | <0.001 |
| **Log<sub>10</sub> Pop Density (1/sq. miles)** | 0.6712 | 0.029 | <0.001 |
| **% Female** | 0.0484 | 0.006 | <0.001 |
| **% Black** | -0.0083 | 0.001 | <0.001 |
| **% Hispanic** | 0.0007 | 0.002 | 0.64 |
| **% Asian** | -0.0095 | 0.002 | <0.001 |
| **% Other** | -0.0262 | 0.004 | <0.001 |
| **% 18 to 40** | -0.0188 | 0.005 | <0.001 |
| **% 40 to 60** | 0.084 | 0.007 | <0.001 |
| **% 60 to 80** | 0.0021 | 0.006 | 0.71 |
| **% 80+** | 0.0551 | 0.012 | <0.001 |
| **% \$15k to \$30k** | -0.0161 | 0.007 | 0.01 |
| **% \$30k to \$45k** | -0.0091 | 0.006 | 0.16 |
| **% \$45k to \$60k** | 0.002 | 0.007 | 0.78 |
| **% \$60k to \$100k** | 0.0073 | 0.005 | 0.15 |
| **% \$100k to \$125k** | 0.0325 | 0.009 | <0.001 |
| **% \$125k to \$200k** | 0.0161 | 0.006 | 0.01 |
| **% \$200k+** | 0.0714 | 0.007 | <0.001 |
| **% Unemployed** | -0.0009 | 0.007 | 0.89 |
| **% No health insurance** | -0.0298 | 0.004 | <0.001 |
| **% High School** | 0.038 | 0.005 | <0.001 |
| **% Some College** | -0.0101 | 0.004 | 0.01 |
| **% College** | 0.0713 | 0.005 | <0.001 |
| **% Graduate School** | -0.0207 | 0.005 | <0.001 |
| **% Houses owner-occupied** | -0.0027 | 0.002 | 0.14 |
| **Median House Price (millions of \$)** | -0.00001 | 0.00002 | 0.43 |
| **Alabama** | 0.0001 | 0.003 | 0.98 |
| **Alaska** | -0.0255 | 0.003 | <0.001 |
| **Arizona** | 0.0395 | 0.003 | <0.001 |
| **Arkansas** | 0.0136 | 0.003 | <0.001 |
| **California** | -0.0096 | 0.003 | <0.001 |
| **Colorado** | 0.0501 | 0.003 | <0.001 |
| **Connecticut** | -0.0044 | 0.003 | 0.18 |
| **Delaware** | -0.0278 | 0.003 | <0.001 |
| **District of Columbia** | 0.0121 | 0.005 | 0.01 |
| **Florida** | 0.0235 | 0.003 | <0.001 |
| **Georgia** | 0.0259 | 0.003 | <0.001 |
| **Hawaii** | -0.0315 | 0.003 | <0.001 |
| **Idaho** | -0.0072 | 0.003 | 0.02 |
| **Illinois** | 0.0001 | 0.003 | 0.97 |
| **Indiana** | 0.0114 | 0.003 | <0.001 |
| **Iowa** | 0.0083 | 0.003 | 0.01 |
| **Kansas** | -0.0014 | 0.003 | 0.67 |
| **Kentucky** | -0.0067 | 0.003 | 0.03 |
| **Louisiana** | 0.0012 | 0.003 | 0.69 |
| **Maine** | -0.0189 | 0.003 | <0.001 |
| **Maryland** | -0.0046 | 0.003 | 0.14 |
| **Massachusetts** | -0.0334 | 0.003 | <0.001 |
| **Michigan** | -0.0275 | 0.003 | <0.001 |
| **Minnesota** | 0.036 | 0.003 | <0.001 |
| **Mississippi** | -0.0025 | 0.003 | 0.42 |
| **Missouri** | 0.031 | 0.003 | <0.001 |
| **Montana** | -0.0207 | 0.003 | <0.001 |
| **Nebraska** | 0.0544 | 0.004 | <0.001 |
| **Nevada** | 0.0006 | 0.003 | 0.85 |
| **New Hampshire** | -0.0222 | 0.003 | <0.001 |
| **New Jersey** | -0.0256 | 0.003 | <0.001 |
| **New Mexico** | 0.0001 | 0.003 | 0.97 |
| **New York** | -0.0254 | 0.003 | <0.001 |
| **North Carolina** | 0.0223 | 0.003 | <0.001 |
| **North Dakota** | 0.0236 | 0.005 | <0.001 |
| **Ohio** | -0.005 | 0.003 | 0.09 |
| **Oklahoma** | 0.0094 | 0.003 | <0.001 |
| **Oregon** | -0.0031 | 0.003 | 0.32 |
| **Pennsylvania** | -0.0303 | 0.003 | <0.001 |
| **Rhode Island** | 0.0216 | 0.004 | <0.001 |
| **South Carolina** | 0.0105 | 0.003 | <0.001 |
| **South Dakota** | 0.0069 | 0.004 | 0.09 |
| **Tennessee** | -0.0034 | 0.003 | 0.26 |
| **Texas** | 0.0235 | 0.003 | <0.001 |
| **Utah** | 0.0318 | 0.003 | <0.001 |
| **Vermont** | -0.0224 | 0.004 | <0.001 |
| **Virginia** | -0.0113 | 0.003 | <0.001 |
| **Washington** | -0.0052 | 0.003 | 0.08 |
| **West Virginia** | -0.0213 | 0.003 | <0.001 |
| **Wisconsin** | 0.0155 | 0.003 | <0.001 |

## References

1. Research C for DE and. Real-World Data: Assessing Electronic Health Records and Medical Claims Data To Support Regulatory Decision-Making for Drug and Biological Products. U.S. Food and Drug Administration. Published December 10, 2021. Accessed August 23, 2022. https://www.fda.gov/regulatory-information/search-fda-guidance-documents/real-world-data-assessing-electronic-health-records-and-medical-claims-data-support-regulatory

2. Bykov K, He M, Gagne JJ. Trends in Utilization of Prescribed Controlled Substances in US Commercially Insured Adults, 2004-2019. JAMA Internal Medicine. 2020;180(7):1006–1008. doi:10.1001/jamainternmed.2020.0989

3. Bangerter LR, Clark CN, Jarvis MS, et al. Medication Fill Patterns for Cognitive and Behavioral or Psychological Symptoms of Alzheimer Disease and Related Dementias. JAMA Network Open. 2022;5(6):e2215678. doi:10.1001/jamanetworkopen.2022.15678

4. Eberly LA, Garg L, Yang L, et al. Racial/Ethnic and Socioeconomic Disparities in Management of Incident Paroxysmal Atrial Fibrillation. JAMA Netw Open. 2021;4(2):e210247. doi:10.1001/jamanetworkopen.2021.0247

5. Hotez PJ. “Neglected infections of poverty in the United States of America”. PLoS Negl Trop Dis. 2008 Jun 25;12(6):e256. doi: 10.1371/journal.pntd.0000256. PMID: 18575621; PMCID: PMC2430531.

6. Dickman SL, Himmelstein DU, Woolhandler S. “Inequality and the health-care system in the USA”. The Lancet 2017 Apr 8–14;389(10077);1431–1441. doi: https://doi.org/10.1016/S0140-6736(17)30398-7.

7. Degtiar, I., & Rose, S. (2021). A review of generalizability and transportability. arXiv preprint arXiv:2102.11904.

8. Dahabreh, I. J., Robertson, S. E., Steingrimsson, J. A., Stuart, E. A., & Hernan, M. A. (2020). Extending inferences from a randomized trial to a new target population. Statistics in medicine, 39(14), 1999–2014.

9. Konrad R, Zhang W, Bjarndóttir M, Proaño R. Key considerations when using health insurance claims data in advanced data analyses: an experience report. Health Syst (Basingstoke). 9(4):317–325. doi:10.1080/20476965.2019.1581433

